# Targeting hypoxia-induced circSTT3A decreases breast cancer stem cell formation via degradation of PGK1 protein and serine synthesis

**DOI:** 10.1101/2023.04.28.538664

**Authors:** Ming Xu, Manran Liu, Xinyue Zhou, Yilu Qin, Liping Yang, Siyang Wen, Yuxiang Qiu, Ting Jin, Shangchun Chen, Rui Tang, Yuetong Guo, Yan Sun

**Author notes:** Corresponding author: Yan Sun. Department of Cell Biology and Medical Genetics, Basic Medical School, Chongqing Medical University, #1 Yi-Xue-Yuan Rd., Yu-zhong District, Chongqing 400016, China. Ming Xu, Manran Liu and Xinyue Zhou contributed equally to this work.

## Abstract

**Background:** Hypoxia is a key feature of tumor microenvironment that can cause fundamental changes in cancer cells, and may also lead to the development of breast cancer stem cells (BCSCs) with self-renewal ability. However, the mechanism of hypoxia in inducing BCSCs is not fully understood.

**Methods:** Performing RNA sequence and bioinformatics analysis, a hypoxia specific circular RNA (circRNA), named circSTT3A, was identified in hypoxic breast cancer cells and tissues. The clinical significance of circSTT3A was investigated in breast cancer (BC) tissues and tissue array. The loss and acquisition of circSTT3A were carried out in vivo and in vitro to confirm its functional roles in BCSC stemness maintenance. HIF1α droved circSTT3A expression was evaluated by chromatin immunoprecipitation and dual luciferase reporter assays. RNA pull-down, RNA immunoprecipitation, silver staining detection, mass spectrographic analysis, co-immunoprecipitation assays and western blotting were exerted to determine circSTT3A directly binding with HSP70 and PGK1 proteins. CircSTT3A-mediated serine metabolism was determined by UHPLC-QTRAP-MS system and ELISA kit. BC mouse model was used to assess the effects of circSTT3A/Hsp70/PGK1 on tumorigenesis and chemotherapy resistance in vivo.

**Results:** A novel hypoxia specific circSTT3A was significantly upregulated in clinical breast cancer tissues, and was related to the clinical stage and poor prognosis of BC patients. The hypoxia inducible factor 1 alpha (HIF1α)-regulated circSTT3A has remarkable effect on mammosphere formation in breast cancer cells. Our work revealed that circSTT3A directly interacting with nucleotide-binding domain of heat shock protein 70 (HSP70) increases the ability of HSP70 to recruit phosphoglycerate kinase 1 (PGK1) via its substrate-binding domain, which reduces the ubiquitination of PGK1 and increases the stability of PGK1. The enhanced PGK1 catalyzes 1,3-diphosphoglycerate (1, 3-BPG) into 3-phosphoglycerate (3-PG) leading to 3-PG accumulation and increase of serine synthesis, thus to facilitate BCSC enrichment under hypoxic microenvironment. Loss of circSTT3A or PGK1 substantially induces suppression in tumor initiation and tumor growth, which dramatically increases tumor sensitivity to Doxorubicin treatment in mice.

**Conclusions:** Hypoxia induced circSTT3A/HSP70/PGK1 axis plays a critical role in maintaining BCSC properties and may be meaningful for treating patients with breast cancer.

## Introduction

Breast cancer (BC) is one of the most common malignancies among women threatening women’s health and leading to cancer-related deaths in women around the world [1]. BC patients often receive conventional therapeutic measures, such as surgery, radiotherapy, endocrine therapy, chemotherapy, and immunotherapy, however, it still accounts for 15% of all cancer deaths in women each year [2]. Therefore, exploring the molecular markers and mechanisms in BC is crucial for diagnosis and treatment.

It is established that the hypoxia region of the tumor microenvironment is mainly located in the area that has a rapid proliferation of cancer cells. In the hypoxic environment, hypoxia-inducible factor-1 alpha (HIF1α) is the most critical transcriptional regulator which has been demonstrated to participate in the transcriptome, proteome, and metabolome networks of BC, thus provides advantages for the survival and development of cancer cells [3, 4]. Among them, studies have identified that hypoxia plays a critical role in induction of breast cancer stem cells (BSCS)-like phenotype, characterized by self-renewal and differentiation potential from the BC cells [5]. The CD44 and CD24 as cell surface antigens have been successfully used to isolate BCSC-like populations from BC cell lines and primary tissues. The recent discovery unveiled that non-BCSCs are plastic cell populations, which readily switch from a non-BCSC to BCSC-state. It has been shown tumor cells can convert from a CD44^low^ to a CD44^high^ state in BC. The plasticity is regulated by ZEB1 transcription [6], AMPK-HIF-1α axis [7], HIF2α [8] and so on. However, the mechanism underlying remains poorly understood. It is particularly essential to reveal the regulatory mechanisms of BCSCs formation and maintenance from the perspective of hypoxia in detail, this may optimize existing BC treatment strategies.

As one member of the eukaryotic transcriptome, circular RNAs (circRNAs) have received considerable attention from researchers. CircRNAs are covalently closed endogenous biomolecules with tissue and cell-specific expression patterns [9]. It is reported circRNAs involved in a variety of tumorigenesis and development as disease molecular markers and key regulatory molecules [10]. In recent years, several studies have found that circRNAs are closely associated with hypoxia with multiple functions in the molecular mechanism. For example, circ-0001875 acts as a miR-31-5p sponge to regulate the expression of SP1 which promotes EMT via a TGF-β/Smad2 signal pathway [11]. In addition, HIF-1α-induced circWSB1 has improved the survival and growth of BC cells under hypoxia via destroying the interaction between USP10 and p53 [12]. Furthermore, circHIF1A can regulate the expression of CD44 by sponging miR-580-5p under hypoxic conditions, thereby participating in the formation of BCSCs and promoting the development of BC [13]. These studies suggest that circRNAs play potentially important roles under hypoxic conditions by distinct modes of action at the plasticity of BCSCs. Thus, exploring the molecular mechanism of circRNAs regulating and maintaining the stemness of cells may be an effective way to decipher the occurrence and development of BC.

It is now appreciated that aberrant alterations in enzymes and metabolites are essential for BC cells to adapt to the hypoxic environment [14]. Under hypoxia, metabolic reprogramming of BC cells increases glycolysis and inhibits oxidative phosphorylation [15]. Combination therapies targeting glycolysis enzymes to surmount drug resistance will be a novel viewpoint for cancer treatment regimens. In solid tumors, most enzyme-encoding genes involved in glycolysis, including hexokinase, aldolase, triosephosphate isomerase, glyceraldehyde-3-phosphate dehydrogenase, phosphoglycerate kinase, and pyruvate kinase M1/2, were hyper-activated under hypoxia to provide ATP and intermediary metabolic products for tumorigenesis [16]. Meanwhile, the post-translational modifications of these genes are great significance to examine the potential biological function. Recent studies showed that the tetramer and dimer transition of pyruvate kinase M2 (PKM2) was regulated by ubiquitination, thereby modulating the Warburg effect in the malignant behavior of BC [17]. The ubiquitin-proteasome system-mediated degradation of hexokinase 2 (HK2) was attenuated by the deubiquitinase function of CSN5 to upregulate glycolysis in hepatocellular carcinoma [18]. Therefore, unraveling the post-translational regulation of enzymes will likely provide insights for understanding metabolic reprogramming, and potential targets for combating hypoxia driven BC tumorigenesis.

In this report, we found that a novel circRNA, circSTT3A, is upregulated in BC and plays a key role in maintaining the stemness of BCSC. CircSTT3A interacts with heat shock protein 70 (HSP70) to regulate the stability of phosphoglycerate kinase 1 (PGK1) protein by reducing the ubiquitination of PGK1, which contributes to cellular 3-PG and serine synthesis in maintaining BCSC stemness and tumor chemotherapy resistance. The enhanced circSTT3A serves as a good diagnostic biomarker for breast cancer.

## Materials and methods

### Clinical samples

Human BC tissue and its matched para-carcinoma mammary tissue (a distance of 5 cm at a minimum from the tumor) were collected from BC patients beyond preoperative history of radiotherapy or chemotherapy in the First Affiliated Hospital of Chongqing Medical University. Written notification has been agreed by all patients and this study was ratified by the Ethics Committee of Chongqing Medical University.

### Cell culture

The human BC cells (BT-549, MDA-MB-468, MDA-MB-231, Hs578T, MDA-MB-436, T47D, SKBR3 and MCF-7) were acquired from the American Type Culture Collection (ATCC, USA), and were cultured with RPMI 1640 or DMEM (Gibco, USA), supplemented with 10% fetal bovine serum (Gibco) and 1% streptomycin/ penicillin (Beyotime, Shanghai, China) at 37 °C in incubator with 5% CO2 and 20% O_2_. To create a hypoxic environment, cells were cultured in a tri-gas incubator with 1% O_2_, 94% N2 and 5% CO_2_. To maintain the metabolites effect, cells were cultured in medium supplemented with 2 mM 3-phosphoglycerate (3-PG, Sigma-Aldrich, #P8877) or 2 mM serine (Sigma-Aldrich, #S4500) [19].

### RNA sequencing

Total RNA was isolated from tumor cells exposed to normoxia or hypoxia with TRIzol reagent (Invitrogen). Then RNA with rRNA depletion was subjected to cDNA synthesis and sequencing by Illumina HiSeq sequencer (Lifegenes, Shanghai, China). HTSeq v0.6.1 was used to count reads and identified the differentially expressed genes according to the overlap of reads and gene location. Quantile normalization and differential expression investigation were performed using the R software package.

### Lentivirus, plasmids, siRNAs and cell transfection

The shRNAs particularly against HIF-1α, circSTT3A and their control hairpins were cloned into GV493/GFP lentiviral vector GenePharma (Shanghai, China); the total length of circSTT3A was amplified and inserted into the lentiviral vector pUC57-ciR (Geneseed, Guangzhou, China) to construct circSTT3A expression vector. The tumor cells were infected by lentiviral and screened by puromycin for 14 days. The siRNA against HSP70, PGK1 and negative control were purchased from Geneseed. HSP70 and PGK1 cDNA were amplified and subcloned into pcDNA3.1 (Geneseed). HSP70 and its deletion mutant sequence were severally inserted into pcDNA3.1-Flag plasmid to construct pcDNA3.1-Flag-HSP70 full length (Flag-HSP70-FL), pcDNA3.1-Flag-HSP70-KH1 domain (Flag-HSP70-1), pcDNA3.1-Flag-HSP70-KH2 domain (Flag-HSP70-2), and pcDNA3.1-Flag HSP70-KH3 domain (Flag-HSP70-3) constructure. The plasmids and siRNAs transfected into cells were carried out by lipofectamine 2000 (Invitrogen, Carlsbad, CA, USA). The sequences of shRNA and siRNA were listed in Supplementary Table 1.

### Mammosphere formation assay

Breast CSC-like cells isolated from MDA-MB-231, BT-549, and primary cells of BC tissues were concentrated in six-well plates coated with 2% poly-2-hydroxyethyl methacrylate (Sigma-Aldrich) at a density of 1×10^4^ cells/ml in serum-free DMEM/F12 (Gibco) medium, supplemented with 0.4% albumin from bovine serum (Sigma-Aldrich), 20 ng/ml basic fibroblast growth factor (Invitrogen), 20 ng/ml epidermal growth factor, B27 (Gibco), and 500 mg/ml insulin (Invitrogen). The mammosphere size (diameter>50 µm) and number were calculated to assess the self-renewal capability, and representative picture were captured with the OLYMPUS IX70 microscope (Tokyo, Japan). The formula (amount of mammospheres per well/number of cells seeded per well × 100) was used to compute the mammosphere-forming efficiency (MFE).

### RNA isolation and qRT-PCR

Extracted total RNA with Trizol reagent was on the basis of the manufacturer’s instructions (Takara, Japan) in both tissues and cells, and was reversed into cDNA using PrimeScriptP RT kit (Perfect Real-Time; Takara). Then cDNA and SYBR Premium Ex Taq II were subjected to qRT-PCR reaction. Consequences were normalized to GAPDH or β-actin expression. All assays were executed at least three times. The relative gene expression was analyzed with the 2^−ΔΔCT^ method. The primer sequences used in qRT-PCR were displayed in Supplementary Table 1.

### Actinomycin D treatment and MG132 treatment

BC cells were incubated with 2 μg/ml Actinomycin D (Sigma-Aldrich) for 0 h, 12 h and 24 h individually. The stability of RNA was anatomized by qRT-PCR. MG132 treatment was carried out by incubating cells with 20 μM MG132 (MedChemExpress, USA) for 12 h, and Co-IP and Western blotting were executed to determine protein and its ubiquitination levels.

### RNase R treatment

The stability of circRNA was verified by RNase R treatment test. Total RNAs of BC cells were divided into two groups treated with or without 1 μl RNase R (20 U/μl; Epicenter, WI, USA) at 37 °C for 30 min, then incubated at 70 °C for 10 min to inactivate RNase R. The treated RNA was then reverse-transcribed for RT-PCR experiment.

### Cytosolic/Nuclear fractionation

Separation of nuclear and cytoplasmic RNA was carried out using PARITM Kit (Invitrogen, USA) in accordance with the manufacturer’s instructions. The reverse transcription and qRT-PCR were executed with extracted RNAs immediately to test the target gene expression. U6 RNA or GAPDH mRNA was the nuclear control or cytoplasmic control in the experiments.

### RNA immunoprecipitation (RIP)

RIP experiment was optimized using the Magna RIP RNA-binding protein immunoprecipitation kit (Millipore, USA) in light of the manufacturer’s instructions. Briefly, 2×10^7^ cells were lysed in RIP lysis buffer on ice. Appropriate amount of cell lysate was pretreated with 5 µg of HSP70 antibody for 30 min at room temperature, then incubated with bead antibody complexes in RIP immunoprecipitation buffer overnight at 4 °C. Bead-bound immunoprecipitation was purged using the elution buffer at 55 °C for 30 min. Phenol/chloroform/isoamyl alcohol was used for RNAs isolating and the purified RNA was used in qRT-PCR.

### RNA pull-down

RNA pull-down experiments were conducted with the Pierce™ Magnetic RNA-Protein Pull-Down kit (Termo Fisher Scientifc) depending on the manufacturer’s instructions. To be brief, the sense or antisense biotin-labeled specific probes against circSTT3A was incubated with MDA-MB-231 cell lysate mingled with streptavidin-linked magnetic beads. The purified proteins were analyzed by mass spectrometry and western blotting. All procedures were implemented under RNase-free circumstance.

### Dual-luciferase reporter assay

In view of the experimental devise for HIF-1α binding spots, the 100 bp sequence around the P1, P2, or P3 sites of STT3A promoter was constructed into pGL3-basic luciferase reporter vector (WT). The P1, P2, or P3 mutation sequence were structured into pGL3-basic vector to get the mutant constructors (Mut). Vectors were co-transfected with pcDNA3.1/HIF1a vector into MDA-MB-231 cells by lipofectamine 2000, respectively. After transfection for 36 h, Dual-Glo Luciferase Assay System (Promega, WI, USA) was used to measure the activities of firefly and Renilla luciferase corresponding to the manufacturer’s protocol.

### Chromatin immunoprecipitation assay (ChIP)

Succinctly, in lined with the protocol of Millipore EZ-ChIP kit (Millipore), 1% of protein-DNA mixture was reserved as the input control, and the rest of mixture was combined with anti-HIF1α antibody (Millipore) or normal rabbit IgG (Millipore) overnight at 4 °C. The protein G Sepharose was used to concentrate the immunoprecipitate complexes. The purified genomic DNA fragments were detected by qRT-PCR.

### Fluorescence in situ hybridization (FISH)

BC cells cultured on coverslips were fixed with 4% paraformaldehyde and infiltrated with 0.5% Triton X-100 for 30 minutes, then incubated with a probe against circSTT3A in hybridization buffer (Geneseed, Guangzhou, China) at 37 °C overnight. After washing with PBS, the cells were incubated with anti-HSP70 antibody at 4 °C overnight, then FITC-labeled goat anti-rabbit IgG was added (Bosterbio, China) at 37 °C for 2 h. DAPI was use to locate nucleus. The pictures were reaped by laser scanning confocal microscope.

### Immunofluorescence (IF) staining

Briefly, BC cells seeded on the coverslips were fixed by 4% polyformaldehyde and permeabilized with 0.5% Triton-X 100, and then sealed with 5% BSA. The cells were incubated overnight with mouse derived anti-HSP70 antibody or rabbit derived anti-PGK1 antibady (GeneTex, USA) at 4 °C, then incubated with appropriate secondary antibodies [Alexa Fluor Plus 555-conjugated anti-mouse (Thermo Fisher scientific, USA) or Alexa Fluor 488-conjugated anti-rabbit (Thermo Fisherscientific, USA)]. After stained with DAPI for nuclei, fluorescent images were captured by confocal microscope.

### Tissue microarray (TMA) and In situ hybridization (ISH)

TMA was prepared from 160 paraffin-embedded samples from Beijing Dingguo Changsheng Biotechnology Co., LTD. ISH was used to detect the expression of circSTT3A in BC tissues following to protocol’s instructions. Briefly, TMA was routinely dewaxed in xylene and gradient ethanol, and hybridized with a specific circSTT3A probe (Geneseed, Guangzhou, China). Then denatured biotin-labeled probe was applied to the section in a wet box at 42 °C overnight. After the sections blocked and treated with alkaline phosphatase-labeled strand avidin, color development was carried out using BCIP/NBT working solution, and checked under microscope.

### Immunohistochemistry (IHC) analysis

IHC staining was performed with Paraffin-embedded tissue sections on the basis of the manufacturer’s protocols. Concisely, after dewaxing and hydration, the slices were incubated with 3% hydrogen peroxide to block the endogenous peroxidase by heating in a microwave for antigen retrieval (AR), and then incubated with primary antibodies against PCNA (1:100), KLF4 (1:100), OCT4(1:100), Ki-67 (1:100) at 4 °C overnight and secondary antibody at room temperature for 1 hour. Carefully washing with T-PBS, sections were colored with diaminobidine (DAB) and counter-stained with hematoxylin and images were photographed by eclipse 80i (Tokyo, Japan).

### CCK-8 and Colony formation assays

To analyze cell viability, Cell Counting Kit-8 (CCK-8, #CK04, Dojindo, Tokyo, Japan) was utilized following the manufacturer’s instructions. Using a microplate reader (#E0226, Beyotime) to evaluate Optical density (OD) value at 450 nm. For colony formation assay, TNBC cells were inoculated into a six-well plate with 1000 cells per well and kept in culture for 14 days to obtain cell colonies, cells were fixed with paraformaldehyde (#P0099, Beyotime) and stained with 0.1% crystal violet (#C0121, Beyotime).

### Transwell assays

After transfection, 3×10^4^ TNBC cells suspended in 200 μL in serum-free medium and seeded into the upper chambers (3374, Corning, USA). After incubation for 8-12 h, wipe the cells in the upper cavity with a cotton swab, fix the cells in the lower cavity in 2% paraformaldehyde, and dye with crystal violet. The number of invasive cells was randomly selected from five selected areas under the microscope.

### Flow cytometry

For cell cycle analysis, TNBC cells were resuspended and immobilized with ice cold 70% ethanol overnight at 4 °C, and examined by the flow cytometry (Becon Dickinson FACS Calibur, NY, USA). For CSC-like cells selection, the mammospheres were separated with 0.25% trypsin and re-suspended into single cell in PBS, then combined with APC anti-CD44 (BD) and PE anti-CD24 (BD) for 30 minutes at 4 °C in the dark. The re-suspended tumor cells in 500 µl PBS were used for selecting the CD44^+^/CD24^−/low^ cell populations by flow cytometry.

### Co-immunoprecipitation (Co-IP)

Co-immunoprecipitation was implemented with antibodies against HSP70 (Abcam) or PGK1 (CST), IgG control (CST) and Pierce™ Classic Magnetic IP/Co-IP Kit (Termo Fisher Scientifc). Briefly, cells were enriched and lysed on ice for 20 minutes with IP lysis/washing buffer with protease inhibitor, then centrifugated at 14,000 g for 20 min. After incubating the collected supernatant with antibodies (5 μg) on the rotator at 4 °C overnight, 25 μL Protein A/G magnetic beads were added and incubated with lysate/antibody mixture at 4 °C for 4 hours. After washing the beads with IP cracking/washing buffer and ultrapure water, 100 μL Lane Marker sample buffer were applied to elute protein and heated at 100 °C for 10 minutes, following appraised by western blotting.

### Western blotting

Total proteins were extracted using Cold RIPA lysis buffer and electrophoresed by 8% or 10% SDS-PAGE, and transferred to PVDF membrane (Millipore, Massachusetts, USA). Blocked with 5% skimmed milk, membranes were incubated with primary antibodies described as follow: HSP70 (1:1000, Cell Signaling Technology), PGK1 (1:1000, CST), c-Myc (1:1000, Abcam), CD44 (1:400, Abcam), KLF4 (1:1000, Millipore), β-actin (1:5000, Abcam). After incubated with HRP-conjugated secondary antibody (1: 3000, Beyotime, Shanghai, China), immunoreactive bands were visualized using ECL detection reagents (Bio-rad, America) on the enhanced chemiluminescence system (Amersham Pharmacia Biotech).

### Metabonomics analysis and Serine measurement

Three replicates from each treatment conditions containing 1×10^7^ cells in 10 cm dishes were collected. After rapid freezing in liquid nitrogen, the cells were temporarily stored at -80 °C, and metabolomics analysis was performed using an UHPLC (1290 Infinity LC, Agilent Technologies) coupled to a QTRAP MS (6500+, Sciex) in Shanghai Applied Protein Technology Co., Ltd. All experiments were completed at three times. The serine concentration of tumor tissues was detected using ELISA kit (Huabbio, China), following the manufacturer’s instructions. The absorbance at 450 nm was measured by microplate reader (SpectraMax M2, Molecular Devices).

### Xenograft mouse model

Animal experiments were performed according to guidelines on animal care approved by the Chongqing Medical University Experimental Animal Management Committee. For tumor initiation assay, different quantity gradients (1×10^3^, 10^4^, 10^5^) of tumor cells in 100 µl of PBS/Matrigel mixture at a 1:1 ratio and were injected into one inguinal mammary fat pad of 5-week-old female nude mice (8 mice per group) subcutaneously. Tumor-initiating potential was evaluated as the ability to form a palpable tumor mass > 0.1 mm^3^ volume. Frequency of stem cells was measured in the light of the Extreme Limiting Dilution Analysis (http://bioinf.wehi.edu. au/software/elda/). For the tumor growth experiment, MDA-MB-231 mammosphere cells (1×10^5^) of the corresponding groups were inoculated into the mammary fat pad of female nude mice (8 mice in each group). The mice with exogenous 3-PG supplementation were treated by intraperitoneal injection of 3-PG at the dose of 100 mg/kg daily for a week prior to cell injection and the whole time until the termination of the animal study. When the tumor volume was about 100 mm^3^, 4 mg/kg doxorubicin was intraperitoneally injected every 5 days. Mice in control group were treated with sterile normal saline (group 1). The xenografts were measured every 5 days with Digital caliper, tumor volume calculation formula is: volume = (length × width^2^)/2. Six weeks after injection, the mice were euthanized. The tumors were isolated, weighed and pictured. The tumor tissues were fixed with 10% formalin, embedded with PARAFN, and cut into 4 μm sections for immunohistochemical analysis.

### Statistical analysis

The whole statistical analyses were implemented with GraphPad Prism 8.0 (GraphPad Software, San Diego, CA). The data were expressed as mean ± standard deviation (SD). As shown in, use student t-test or one-way ANOVA for statistical comparison. P<0.05 was considered statistically significant.

## Results

### CircSTT3A is a hypoxia induced circRNA in breast cancer

Hypoxia is a universal feature in the microenvironment of solid tumors. To understand the potential circRNA involved in regulating biological features of BC under hypoxic environment, RNA-seq analysis was used to identify circRNAs expression profiles of MDA-MB-231 cells exposed to normoxia (Norx, 20% O_2_) or hypoxia (Hypx, 1% O_2_) for 24 h. 247 upregulated and 197 downregulated circRNAs were distinguished (log_2_ |Fold change| ≥2, FDR <0.05) and the top 50 differently upregulated circRNAs were depicted by the heatmap (Fig. 1a). Simultaneously, we performed bioinformatics analysis on the microarray profiles (GSE101124) of BC tissues from the Gene Expression Omnibus (GEO) database, and 46 upregulated circRNAs were ascertained (log_2_ |Fold change| ≥1, FDR <0.05). The intersections of circRNAs between our RNA-seq profiles and GSE101124 were obtained by Venny 2.1.0, in which circSTT3A (hsa_circ_0024760), circSTIL (hsa_circ_0002632) and circCDC25A (hsa_circ_0002023) were the highestly expressed circRNAs related to hypoxia and BC (Fig. 1b). Since circSTT3A was the most significantly increased one in MDA-MB-231 cells under hypoxia, which was finally chosen for the further studies (Supplementary Fig. 1a). Then, the enhanced hypoxic circSTT3A was further confirmed by qRT-PCR in a series of hypoxic BC cell lines (Fig. 1c). The circSTT3A (996 nt) is a circular transcript from the back-splicing of the exons 2-9 of STT3A gene located on human chromosome 11q24.2 (Fig. 1d). Subsequently, the covalent closed-loop structure of endogenous circSTT3A was determined by RT-PCR with convergent and divergent primers, along with GAPDH as a negative control (Fig. 1e). It was found that circSTT3A could be augmented by divergent primers in complementary DNA (cDNA) rather than in genomic DNA (gDNA). Furthermore, as shown in Fig. 1f-1g, circSTT3A was resisted to the digestion of RNase R, while both linear STT3A and GAPDH mRNA were dramatically degraded. Similarly, treatment with actinomycin D demonstrated that the stability of circSTT3A was much better than linear STT3A (Fig. 1h). Moreover, both cytoplasmic/nuclear fractional experiments and RNA fluorescence in situ hybridization (FISH) revealed that circSTT3A was mainly located in the cytoplasm of BC cells in normoxia and hypoxia condition (Fig. 1i-1j). These results suggest that circSTT3A is a hypoxia-related circRNA and mainly locates in the cytoplasm of BC cells.

**Figure 1.**
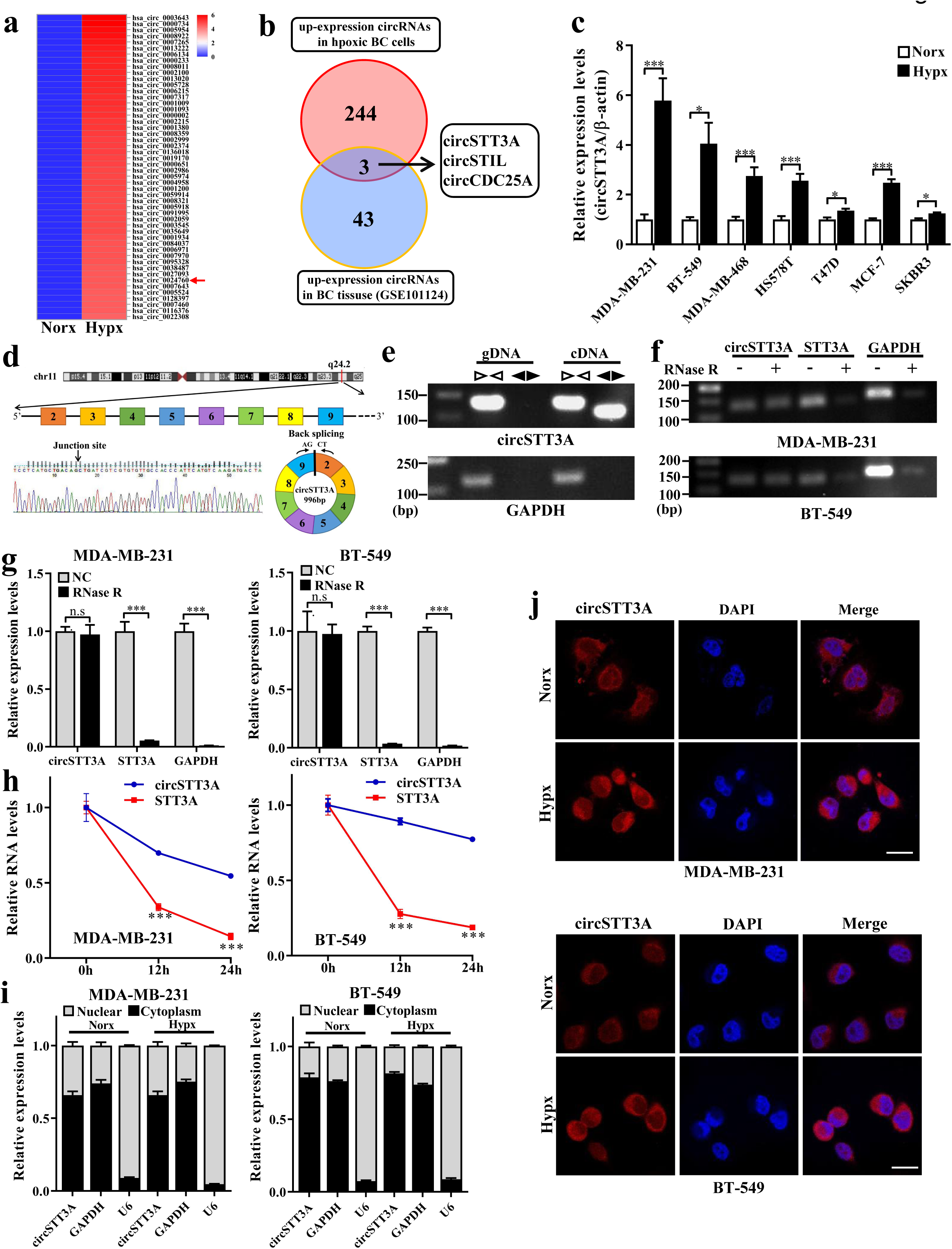
Profiling of dysregulated circRNAs and aberrant expression of circSTT3A in hypoxic BC cells. **a.** Heatmap to show the top 50 differently upregulated circRNAs of MDA-MB-231 exposed to hypoxia (Hypx, 1% O_2_) compared with normoxia (Norx, 20% O_2_) for 24 h. **b.** Venn diagram shows the circSTT3A, circSTIL and circCDC25A were the major up-expressed circRNAs identified by our microarray chip data and in BC tissues microarray data (GSE101124). **c.** qRT-PCR was used to detect the relative expression of circSTT3A in multiple BC cell lines under normoxia and hypoxia for 24 h. **d.** Schematic representation of circSTT3A generated by host gene STT3A. **e.** Gel electrophoresis showed circSTT3A and GAPDH were amplified from MDA-MB-231 gDNA and cDNA using convergent and divergent primers. **f-g.** PCR **(f)** and qRT-PCR **(g)** were used to detect the abundances of circSTT3A and linear STT3A mRNA in BC cells after RNaseR treatment. **h.** qRT-PCR was used to detect the expression of circSTT3A and STT3A in BC cells treated with actinomycin D. **i-j.** The subcellular localization of circSTT3A in MDA-MB-231 and BT-549 cells cultured in hypoxia and normoxia was determined by Nuclear-cytoplasmic fractional assay **(i)** and FISH **(j)** (Scale bar, 25 μm). Data were presented as the mean±SD, *p<0.05, ***p < 0.001.

### Hypoxic circSTT3A is upregulated by HIF-1α

It is established that HIF-1α is a key hypoxia-related transcription factor, which can modulate the expression of various hypoxia-related genes. In compare with control BC cells, loss of HIF-1α led to dramatically down-regulated circSTT3A in the hypoxic MDA-MB-231cells identified by the RNA-seq (Fig. 2a), suggesting that HIF-1α facilitates the expression of circSTT3A in BC cells under hypoxia. Furthermore, analyzed by bioinformatics using Promo database and Ominer database, we identified 7 overlapping transcription factors (TFs), which included HIF-1α, the hypoxia specific TF, potentially transcript its mother gene STT3A (Fig. 2b). For further verify HIF-1α is the regulator for circSTT3A, these TFs were silenced by small interfering RNAs (siRNA) in MDA-MB-231 cells under hypoxia and circSTT3A levels were checked (Supplementary Fig. 1b-1c). HIF-1α knockdown rather than other TFs resulted in a conspicuous reduction of circSTT3A under hypoxia (Fig. 2c, Supplementary Fig. 1c-1d). Inversely, lentivirus-carried ectopic HIF-1α significantly increased circSTT3A expression in normoxic MDA-MB-231 cells (Fig. 2d, Supplementary Fig. 1e). These findings confirmed that circSTT3A expression was dependent on HIF-1α. Subsequently, JASPAR database were used to analyze the binding motifs of STT3A promoter for HIF-1α and 3 binding sites (scores > 9) were selected for further study. The 3 binding sites of the wild type (WT) or mutant (Mut) STT3A sequence were constructed into luciferase reporter vector, respectively (Fig. 2e). The luciferase activity revealed that the P2 and P3 WT sites were the potential HIF-1α binding sites in hypoxic MDA-MB-231 cells (Fig. 2f) or in nornoxic MDA-MB-231 cells with ectopic HIF-1α (Fig. 2g). Moreover, HIF-1α binding to the P2 and P3 sites of the STT3A promoter was further verified by ChIP-qPCR analysis (Fig. 2h). Taken together, HIF-1α drives circSTT3A expression under hypoxia conditions.

**Figure 2.**
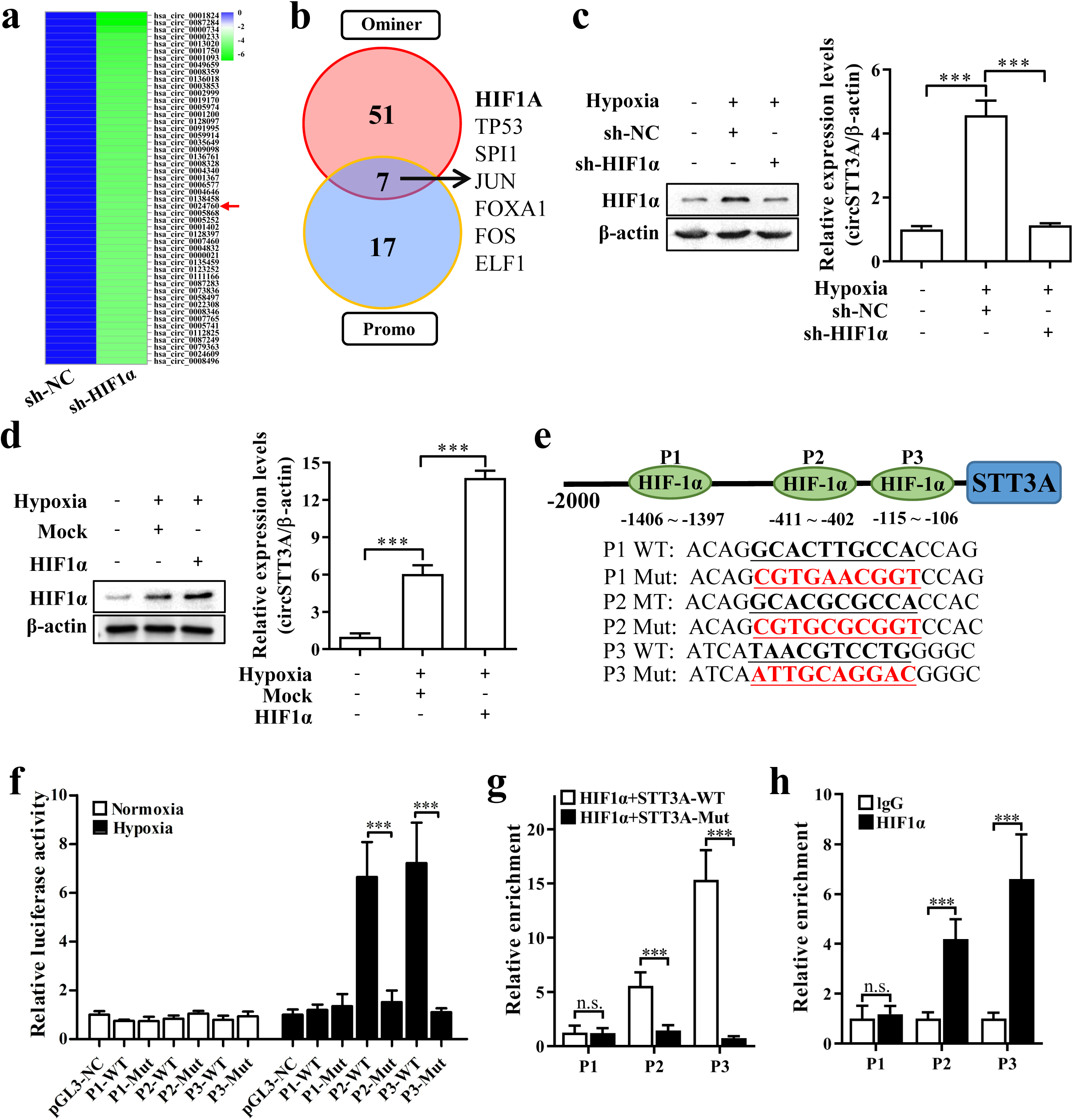
HIF-1α binding to STT3A promoter enhances circSTT3A expression under hypoxia. **a.** Heatmap denoted the top 50 downregulated circRNAs of MDA-MB-231 after knockdown of HIF1α. **b.** Venn plot represented seven potential transcription factors of STT3A promoter in Promo and Ominer datasets. **c-d.** Western blotting and qRT-PCR were used to analyze the HIF1α protein level and the circSTT3A expression in BC cells after knockdown **(c)** or over-expression **(d)** of HIF1α. **e.** Schematic illustration revealed three predicted positions of HIF1α and the wild type (WT) and mutant (Mut) sequences in -2000bp STT3A promoter. **f-g.** The relative luciferase activities were detected in MDA-MB-231 cells transfected with reporter plasmid containing wild type or mutant STT3A promoter sequence in different oxygen content conditions **(f)** and over-expression of HIF-1α **(g)**. **h.** The potential binding sites of HIF-1α and STT3A promoter in MDA-MB-231 cells were detected by ChIP-qPCR. IgG was used as a negative control. Data were presented as the mean ± SD, ***p < 0.001.

### The enhanced circSTT3A is correlated with BC patient progression and works as biomarker of BC

To investigate the clinical significance of circSTT3A in BC, we performed qRT-PCR assays in 60 pairs of clinical specimens of BC tissues and paracancerous tissues. The result revealed that the expression of circSTT3A was observably increased in tumor tissues (Fig. 3a-3b). Moreover, the ROC analysis clarified that the expression of circSTT3A had a meaningful diagnostic value (AUC=0.738, P<0.0001) on BC patients (Fig. 3c). Subsequently, ISH was applied to detect the expression of circSTT3A on 160 BC TMAs, the representative IHC images were displayed in Figure 3d, and the enhanced circSTT3A in BC tissues was closely associated with histology grade (p=0.038) and advanced T stage (p=0.026) (Fig. 3e-3f and Table 1); Kaplan-Meier survival analysis further revealed that patients with high level of circSTT3A were significantly related to poor prognosis (Fig. 3g). In addition, equivalent results also occurred in the histological grade II-III and T stage I-II (Fig. 3h-3i). These results highlight that the circSTT3A acts as an oncogenic circRNA and is potentially correlated with BC progression.

**Figure 3.**
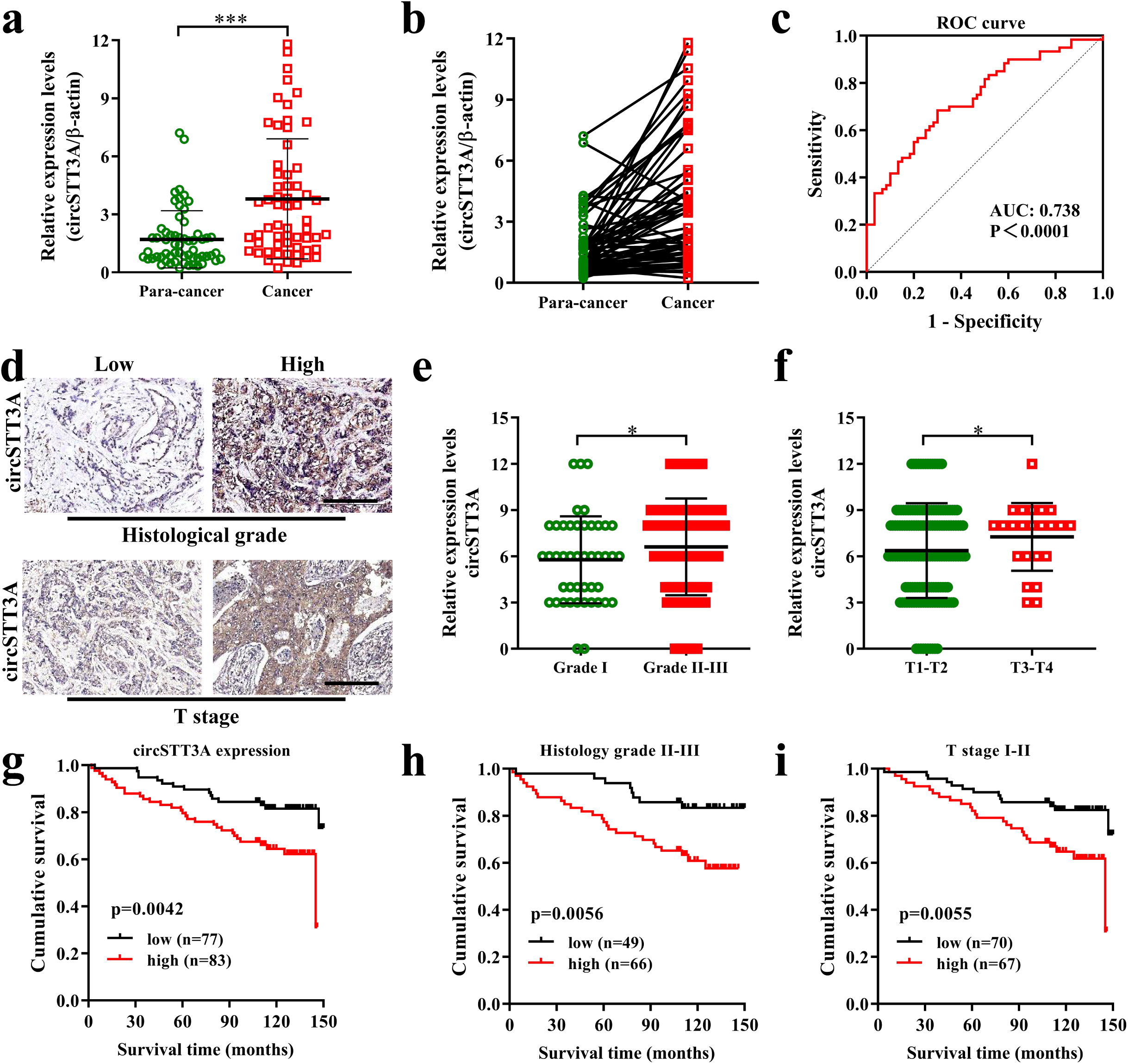
CircSTT3A is up-regulated in BC and related to the progression and poor prognosis of BC patients. **a-b.** Relative expression of circSTT3A in 60 pairs of BC and adjacent tissues. **c.** ROC curve was used to estimate the diagnostic value of circSTT3A for BC. **d.** Representative ISH staining images of circSTT3A on TMAs (Scale bar, 200 μm). **e-f.** Quantification of ISH score of circSTT3A in BC pathological grade and stage. **g-i.** Kaplan–Meier survival curves of BC patients according to the circSTT3A level. Data were presented as the mean ± SD, *p<0.05, ***p < 0.001.

**Table 1.** Correlation between circSTT3A expression and clinicopathological features in 160 BC patients

| Clinicopathological features | Group | circSTT3A |  | *P* value |
| --- | --- | --- | --- | --- |
|  |  | low | high |  |
| Age (years) | <50 | 37 | 34 | 0.369 |
|  | ≥50 | 40 | 49 |  |
| Histology grade | I | 28 | 17 | 0.026 |
|  | II–III | 49 | 66 |  |
| T stage | 1-2 | 70 | 67 | 0.038 |
|  | 3-4 | 7 | 16 |  |
| N stage | N0 | 32 | 33 | 0.817 |
|  | N1-N3 | 45 | 50 |  |
| TNM stage | I–II | 53 | 46 | 0.082 |
|  | III | 24 | 37 |  |

### CircSTT3A plays a notable role in maintaining BCSCs stemness

To explore the oncogenic role of circSTT3A in BC cells, cell models with stable knockdown or overexpression of circSTT3A were constructed (Supplementary Fig. 1f-1g). It is noteworthy that ectopic overexpression or loss of circSTT3A had no effect on the expression of STT3A mRNA (Supplementary Fig. 1f-1g). Next, we tried to examine the potential biological function of circSTT3A via cell viability assay, flow cytometry analysis, invasion assays, and mammosphere formation assay (Supplementary Fig. 2a-2c and Fig. 4a-4b). Interestingly, the results revealed that circSTT3A had remarkable effect on mammosphere formation in BC cells (Fig. 4a-4b). The forming efficiency of mammospheres was evidently lessened in circSTT3A knockdown BC cells under hypoxia condition and meanwhile strengthened in BC cells with ectopic circSTT3A overexpression under normoxia environment, which suggested that circSTT3A may involve in regulating the function of breast cancer stem cells (BCSCs) (Fig. 4a-4b). According to these findings, the mRNA and proteins levels of stemness-related markers (CD44, c-Myc and Klf4) were evaluated and the results exhibited the same trend in these cells (Fig. 4c and Supplementary Fig. 2d). To validate whether the similar phenomenon be existed in primary BC cells, we isolated the primary cells from BC tumor and certified that the primary BC cells with high circSTT3A expression had higher mammosphere-formation abilities than these with low circSTT3A (Fig. 4d). Further analysis by flow cytometry indicated that mammospheres derived from the high circSTT3A level of primary BC cells appeared higher percentage in CD44^+^/CD24^−/low^ cell populations in comparison to those derived from BC cells with low circSTT3A expression (Fig. 4e). Due to BCSCs were considered to be the major cause of BC initiation, we explored whether circSTT3A could contribute to BCSCs-derived tumor initiation by limiting dilution analysis in vivo (Fig. 4f-4g). The results demonstrated that over-expression of ectopic circSTT3A in normoxic MDA-MB-231 cells significantly increased the tumorigenesis of spheroid cells. On the contrary, the tumor-initiating capacity was impaired by the circSTT3A knockdown in hypoxic MDA-MB-231. Furthermore, we measured the volumes of xenograft tumors which injected with 1×10^5^ of indicated cells, the results certified repressive circSTT3A should be beneficial to tumor growth (Fig. 4h). Similarly, the primary BC cells with high-expressed circSTT3A also had stronger tumor-initiating, tumorigenesis and tumor growth abilities than the low circSTT3A expression ones (Fig. 4i-4j). All together, these evidences indicate that circSTT3A exerts positive influence on maintaining BCSCs’ stemness and BC development.

**Figure 4.**
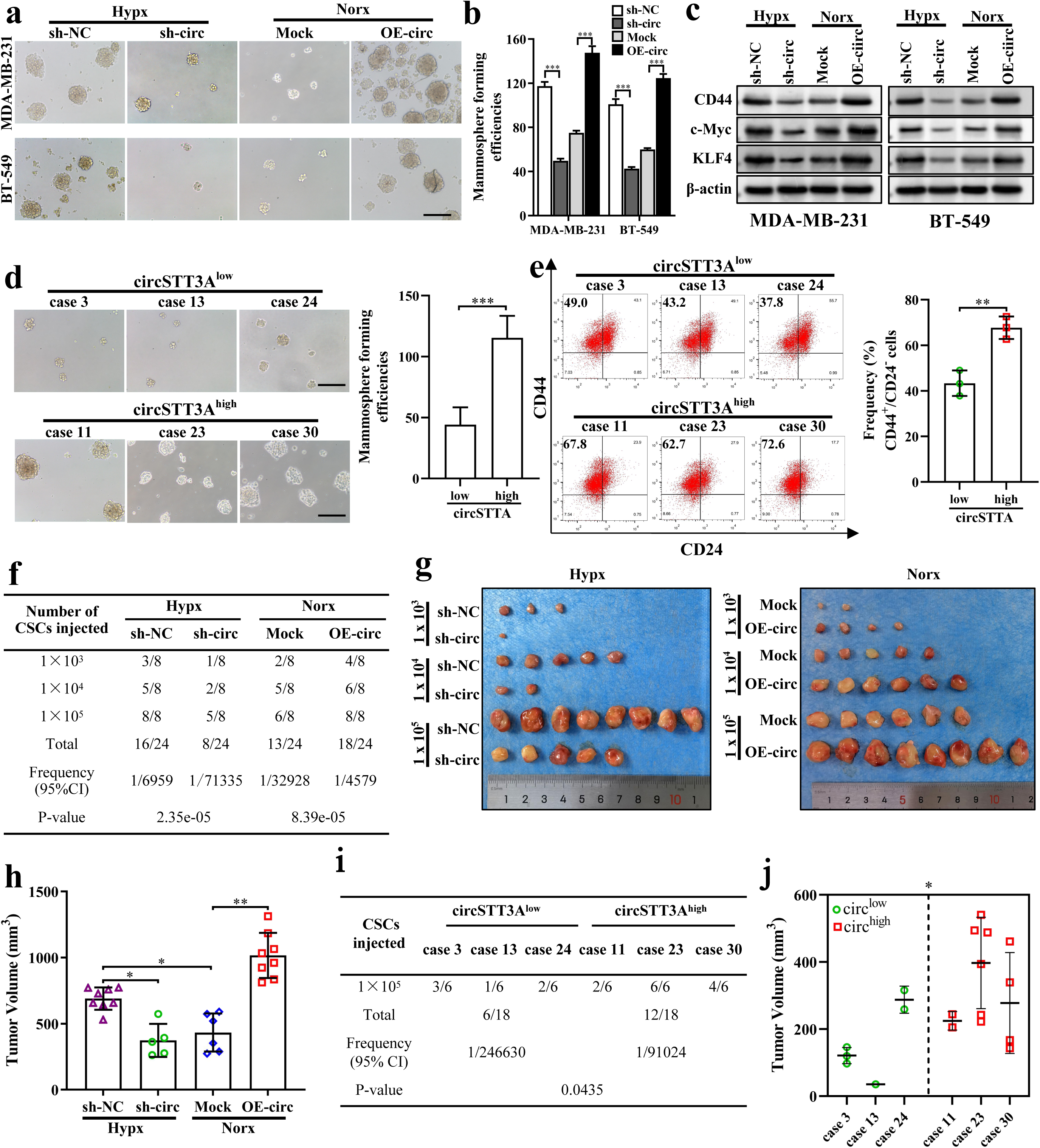
CircSTT3A is required to maintain the stemness of BCSCs. **a-b.** Representative mammosphere images **(a)** and mammosphere formation efficiency **(b)** of the indicated BC cells with ectopic circSTT3A overexpression under normoxia or circSTT3A knockdown under hypoxia (Scale bar, 200 μm). **c.** Western blotting was used to analyze the expression of stemness-related proteins c-Myc, KLF4 and SOX2 in BC cells as mentioned in A and B. **d.** Representative mammosphere images of primary BC cells with low or high circSTT3A (Scale bar, 200 μm). **e.** Flow cytometry was used to determine the percentage of CD44^+^/CD24^-^ cell population in representative BC tissues with low or high circSTT3A. **f.** The MDA-MB-231 derived mammospheres were limit diluted (1×10^3^, 1×10^4^ and 1×10^5^) and subcutaneously implanted into mammary fat pad of nude mice to investigate the effects of circSTT3A on tumor-initiation (n=8 for each mice group). **g-h.** Representative pictures of xenograft tumors **(g)** and xenograft tumor (1×10^5^/mouse) volume **(h)**. **j-k** Mammospheres cultured from BC tissues with low or high circSTT3A (1×10^5^/mouse, n=6 for each mice group) were used to study the effects of circSTT3A on tumor-initiation **(j)** and xenograft tumor (1×10^5^/mouse) volume **(k)**. Data was presented as the mean±SD, *p<0.05, **p<0.01 and ***p < 0.001.

### CircSTT3A directly binding with HSP70 recruits PGK1 and maintains PGK1 stability

To further decipher the latent molecular mechanism of circSTT3A in maintaining the stemness of BC cells, we utilized the RNA-pulldown experiment under hypoxia MDA-MB-231. The silver staining detection, mass spectrographic analysis and western blotting were exerted to determine the circSTT3A-conjugated proteins, and Heat Shock Protein 70 (HSP70) was found to be significantly concentrated on circSTT3A (Fig. 5a), meanwhile the enrichment of circSTT3A on HSP70 was subsequently corroborated by RIP assay (Fig. 5b). The circSTT3A-mediated HSP70 enrichment mainly located in cytoplasm of hypoxic BC cells (Fig. 5c). In the meantime, we appraised the effect of circSTT3A knockdown under hypoxia or ectopic circSTT3A under normoxia on HSP70 expression, qRT-PCR and western blotting experiments revealed that the HSP70 level had no significant fluctuation to circSTT3A status (Fig. 5d-5e). In order to determine the HSP70-interacting specific fragment of circSTT3A, we investigated the HSP70 function domains by UniProt databese and constructed Flag-tagged full-length and three truncated fragments of HSP70 (Fig. 5f). Using RNA pull down and RIP assays, we revealed that the first fragment of HSP70 (Flag-HSP70-1, Nucleotide-binding domain, NBD, 1-386aa) was crucial for its interaction with circSTT3A directly (Fig. 5g-5h). These data show that circSTT3A should recruit HSP70 through directly binding with HSP70-NBD, but not involve in regulating HSP70 transcription.

**Figure 5.**
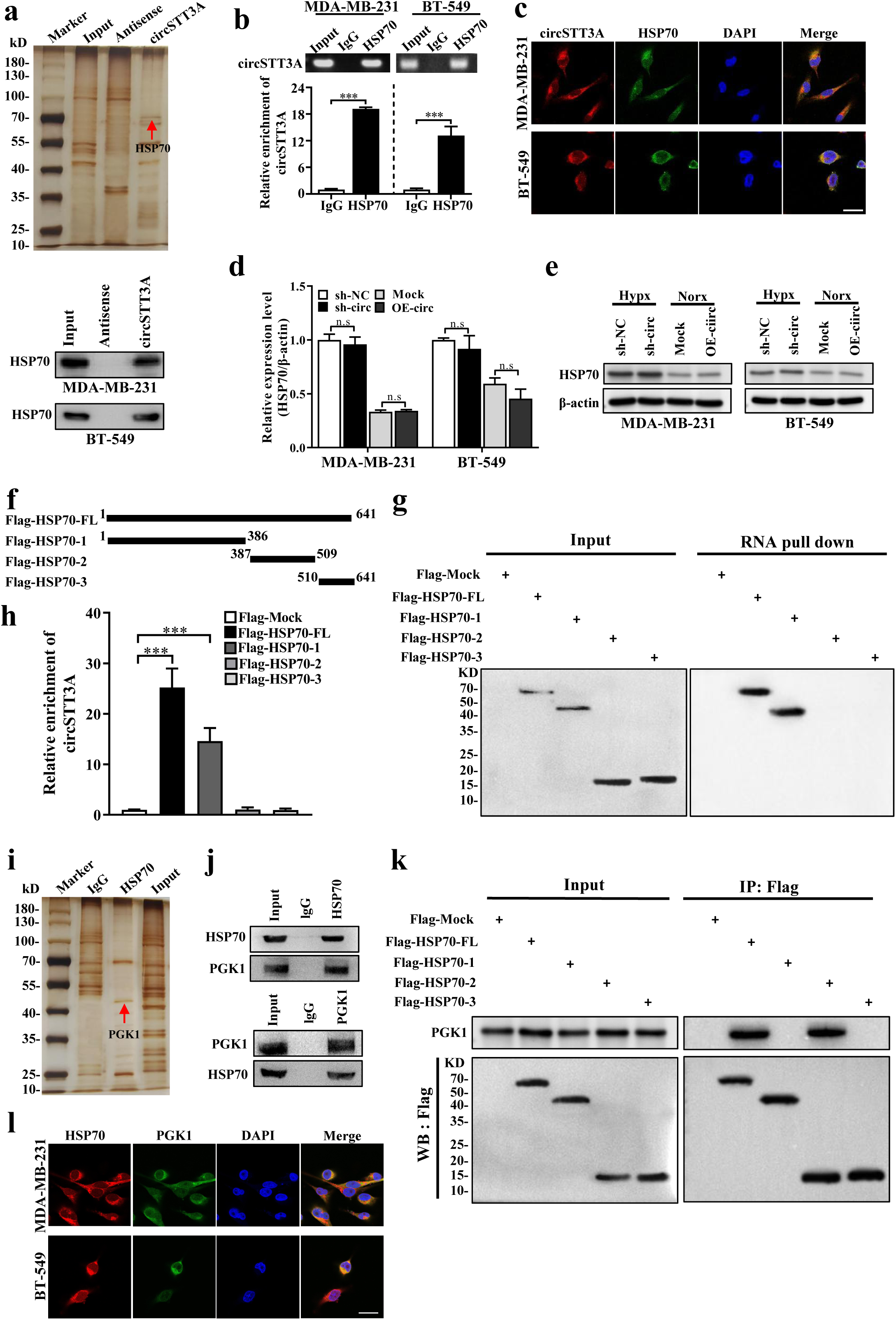
CircSTT3A binding with HSP70 promotes recruitment of PGK1. **a.** Silver staining of proteins pulled down by circSTT3a-specific biotin-labeled probe and control probe. HSP70 was ascertained as a candidate protein interacting with circSTT3A by RNA pull-down. **b.** The interaction between circSTT3A and HSP70 was verified by RIP and qRT-PCR in MDA-MB-231 and BT-549 cell lysate. **c.** FISH and IF co-staining indicated the co-localization of circSTT3A (red) and HSP70 (green) in MDA-MB-231 and BT-549 cells cultured in hypoxia (Scale bar, 25 μm). **d-e**. qRT-PCR and Western blotting were used to detect the effects of hypoxia-induced circSTT3A or ectopic circSTT3A overexpression on the expression of HSP70 in BC cells. **f.** Schematic diagram of HSP70 full-length protein and truncated protein. **g.** RNA pull-down experiment using biotin-labeled circSTT3A probe in hypoxic MDA-MB-231 cells expressing full-length or truncated HSP70, Western blot revealed the enrichmen of indicated full-length of HSP70 and its deletion mutants. **h.** RIP assays were performed with anti-Flag in hypoxic MDA-MB-231 cells transfected with indicated full-length or truncated HSP70 plasmids. **i.** Co-IP assays were carried out in hypoxic MDA-MB-231 cells under indicated conditions using anti-HSP70 and IgG control, followed by silver staining. **j.** The Co-IP experiment used anti-HSP70 or anti-PGK1 to analyze the direct interaction between HSP70 and PGK1 in MDA-MB-231, respectively. **k.** Co-IP using anti-Flag was applied to identify the interaction between the truncated HSP70 protein and PGK1 in MDA-MB-231 cells transfected with indicated full-length or truncated HSP70 plasmids. **l.** IF co-staining showed the co-localization of HSP70 (red) and PGK1 (green) in BC cells (Scale bar, 25 μm). Data were presented as the mean±SD, ***p<0.001.

It has been documented that HSP70 protects the client proteins from ubiquitinmediated degradation via interaction [20]. We hypothesized that circSTT3A might affect the levels of client proteins by binding HSP70. Therefore, Co-IP experiment, SDS-PAGE, and mass spectrometry analysis were performed with an antibody against HSP70 in ectopic circSTT3A overexpressed MDA-MB-231. The potential protein Phosphoglycerate Kinase 1 (PGK1) was identified in the pull-down precipitate, which was co-precipitated with HSP70 in ectopic circSTT3A overexpressed MDA-MB-231 (Fig. 5i), and the direct interaction between HSP70 and PGK1 was further verified by co-IP and WB in hypoxic MDA-MB-231 cells (Fig. 5j). Using Flag-tagged full-length HSP70 and its truncated mutants in CO-IP assays, we confirmed that the second fragment of HSP70 (Flag-HSP70-2, Substrate-binding domain, SBD, 387-509aa) was radical for PGK1 binding (Fig. 5k). In addition, the co-localization of HSP70 and PGK1 was verified in cytoplasm by IF assay (Fig. 5l). Thus, these data prove that circSTT3A and HSP70 could form a circRNA-protein complex to recruit PGK1.

The aforementioned data revealed circSTT3A directly binding with HSP70 recruited PGK1 to form a triple complex. We wondered whether PGK1 levels could be affected. Firstly, the mRNA expression of PGK1 was not affected by circSTT3A under hypoxia or normoxia, while the protein level of PGK1 had a meaningful alteration in the same conditions (Fig. 6a-6b), suggesting that circSTT3A may partake the post-transcriptional regulation of PGK1. Whereafter, the co-IP/WB consequence displayed that HSP70 could enroll more endogenous PGK1 in ectopic circSTT3A overexpressing BC cells (Fig. 6c, down panel); however, knockdown of hypoxic circSTT3A reduced the protein abundance of PGK1 enriched by HSP70 under hypoxia condition (Fig. 6c, upper panel). To further explore the effect of circSTT3A-HSP70 interaction on PGK1 recruitment, PGK1 proteins were determined under circSTT3A knockdown and HSP70 rescue condition (Supplementary Fig 3a). PGK1 protein levels were reduced under knockdown of circSTT3A, yet could be resumed by ectopic HSP70 (Fig. 6d, left panel). In accordance, silence of HSP70 in BC cells (Supplementary Fig 3b) alleviated ectopic circSTT3A-caused enhance of PGK1 (Fig. 6d, right panel). To implement whether circSTT3A modulated PGK1 protein by HSP70, the PGK1 protein stability was tested. Administrating hypoxic BC cells with the proteasome inhibitor MG-132 prevent circSTT3A silence or HSP70 knockdown-induced PGK1 protein degradation (Fig. 6e). Subsequently, as shown in Figure 6f, the ubiquitination level of PGK1 was increased after knockdown of circSTT3A, which could be attenuated by overexpression of ectopic HSP70. These findings signify that circSTT3A-HSP70 maintains PGK1 stability through alleviating PGK1 ubiquitination.

**Figure 6.**
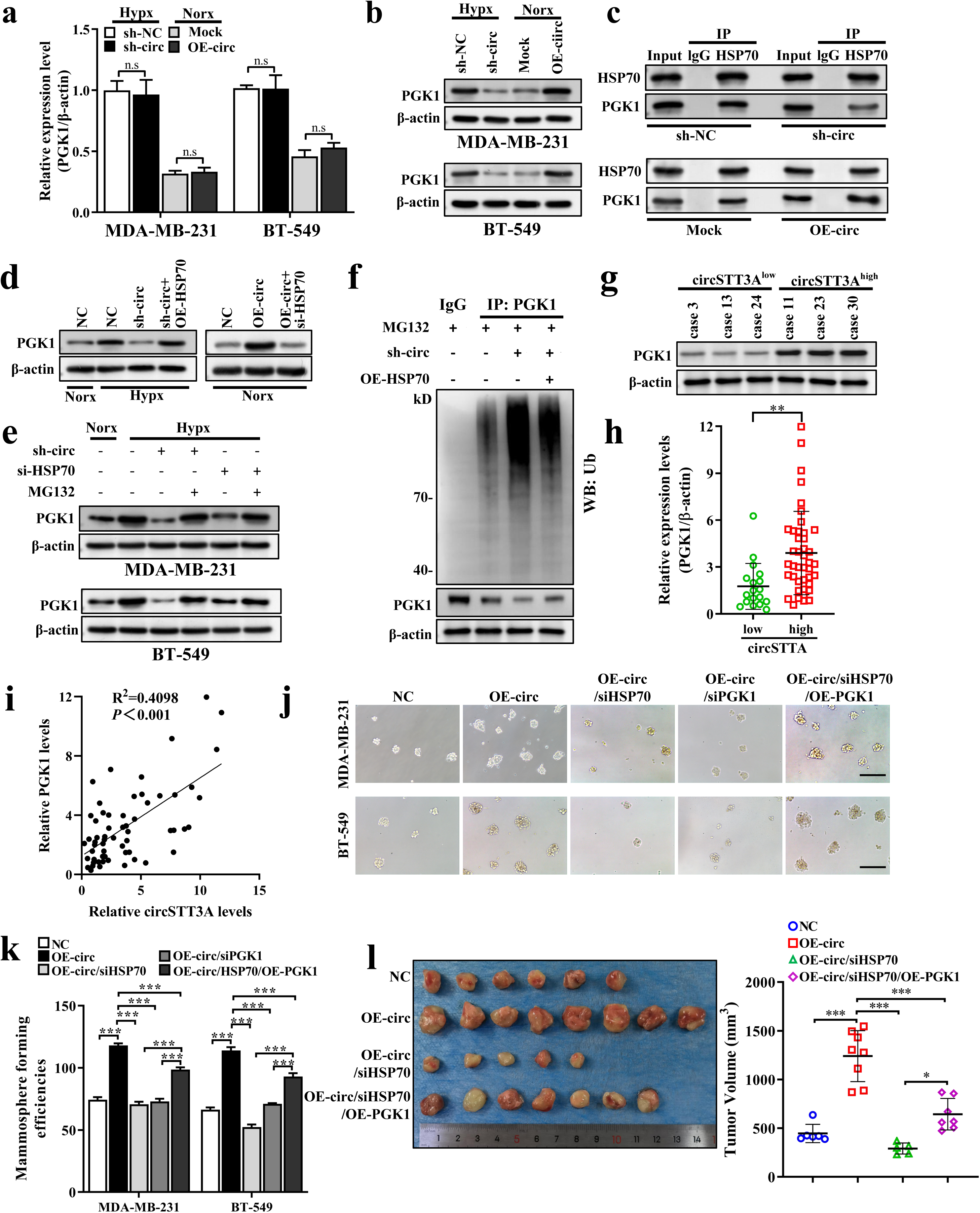
CircSTT3A cooperates with HSP70 to increase PGK1 protein stability and maintained stem cell properties. **a-b.** mRNA and protein levels of PGK1 were determined by qRT-PCR **(a)** and Western blotting **(b)** in BC cells under indicated conditions. **c.** co-IP and WB were carried out in hypoxic MDA-MB-231 cells with or without circSTT3 and in normoxic MDA-MB-231 with or without ectopic circSTT3A expression using anti-HSP70 and anti-PGK1, IgG as negative controls. **d.** The expression of PGK1 was determined after transfection or co-transfection with the indicated vectors or siRNA in MDA-MB-231 cells by Western blotting. **e.** Western blot analysis of PGK1 expression in hypoxic and normoxic BC cells transfected with corresponding siRNA under MG-132 treatment. **f.** Ubiquitination of PGK1 was appraised in MG132-treated MDA-MB-231 cells under indicated conditions. **g-h.** The expression of PGK1 in BC tissues with low or high level of circSTT3A were evaluated by Western blotting **(g)** and qRT-PCR in 60 BC tissues **(h)**. **i.** Correlation analysis of circSTT3A and PGK1 in BC tissues. **j-k.** The representative images **(j)** and quantitative analysis **(k)** of mammosphere formation abilities were showed after transfection or co-transfection with the indicated vectors or siRNA. **l.** The representative images and volume analysis of xenograft tumors for each group (n=8). Data are presented as the mean±SD; *p<0.05, **p<0.01 and ***p < 0.001.

### CircSTT3A-HSP70-induced PGK1 enhance involves in BCSCs maintenance and chemotherapy resistance by regulating PGK1–mediated serine synthesis

To understand whether the enhanced PGK1 involves in circSTT3A-regulated stemness maintenance of BC cells, PGK1 and circSTT3A levels and mammosphere formation were evaluated using tumor tissues and in vitro experiments. The increased PGK1 was perceived in circSTT3A-high breast cancer tissues compared with that in circSTT3A-low tumor tissues (Fig. 6g-6h). Moreover, Pearson correlation analysis illustrated that circSTT3A level was positively correlated with PGK1 expression in BC tumors (Fig. 6i). To study the function of PGK1, ectopic PGK1 overexpression and knockdown BC cells were established (Supplementary Fig 3c-6d). Subsequently, in vitro studies showed that the efficiency of mammosphere formation was elevated in hypoxic BC cells with enhanced circSTT3A, whose mammosphere formation was impaired by loss of HSP70 or PGK1; however, the effect of circSTT3A on mammosphere formation could be recovered by ectopic PGK1 in HSP70-silenced BC cells (Fig. 6j-6k). Furthermore, our in vivo experiments manifested that knockdown of HSP70 in MDA-MB-231/circSTT3A BCSCs significantly decreased tumor initiation and tumor growth, and ectopic PGK1 could offset the absence of HSP70-mediated tumor initiation and tumor growth (Fig. 6l). Taken together, our data demonstrate that circSTT3A binding with HSP70 recruits PGK1 to preserve the stability of PGK1 protein and contributes to BCSC formation and stemness maintenance.

It is well known that PGK1 is a crucial enzyme in the metabolic glycolytic pathway, which catalyzes 1,3-diphosphoglycerate (1,3-BPG) into 3-phosphoglycerate (3-PG), leading to increased serine synthesis via serine synthesis pathway (SSP) [21]. To investigate whether SSP is altered by circSTT3A, we measured the levels of multiple metabolites related with SSP using HPLC-MS. The 3-PG, phosphoserine (pSer) and serine were significantly reduced in hypoxic BC cells when circSTT3A, HSP70 or PGK1 were knocked down, respectively (Fig. 7a). To validate the function of 3-PG and serine on BC cells, exogenous 3-PG and serine were supplemented in the cell culture medium. Addition of 3-PG and serine could notably attenuate the decrease of colony formation (Fig. 7b and Supplementary Fig. 4a), mammosphere formation (Supplementary Fig. 4b) and CSC-related protein reduces (Fig. 7c) caused by loss of circSTT3A, HSP70 or PGK1 in hypoxic BC cells. These data confirmed that circSTT3A-HSP70-PGK1 axis regulates glycolytic pathway leading an enhanced serine, thereby maintaining BCSCs stemness.

**Figure 7.**
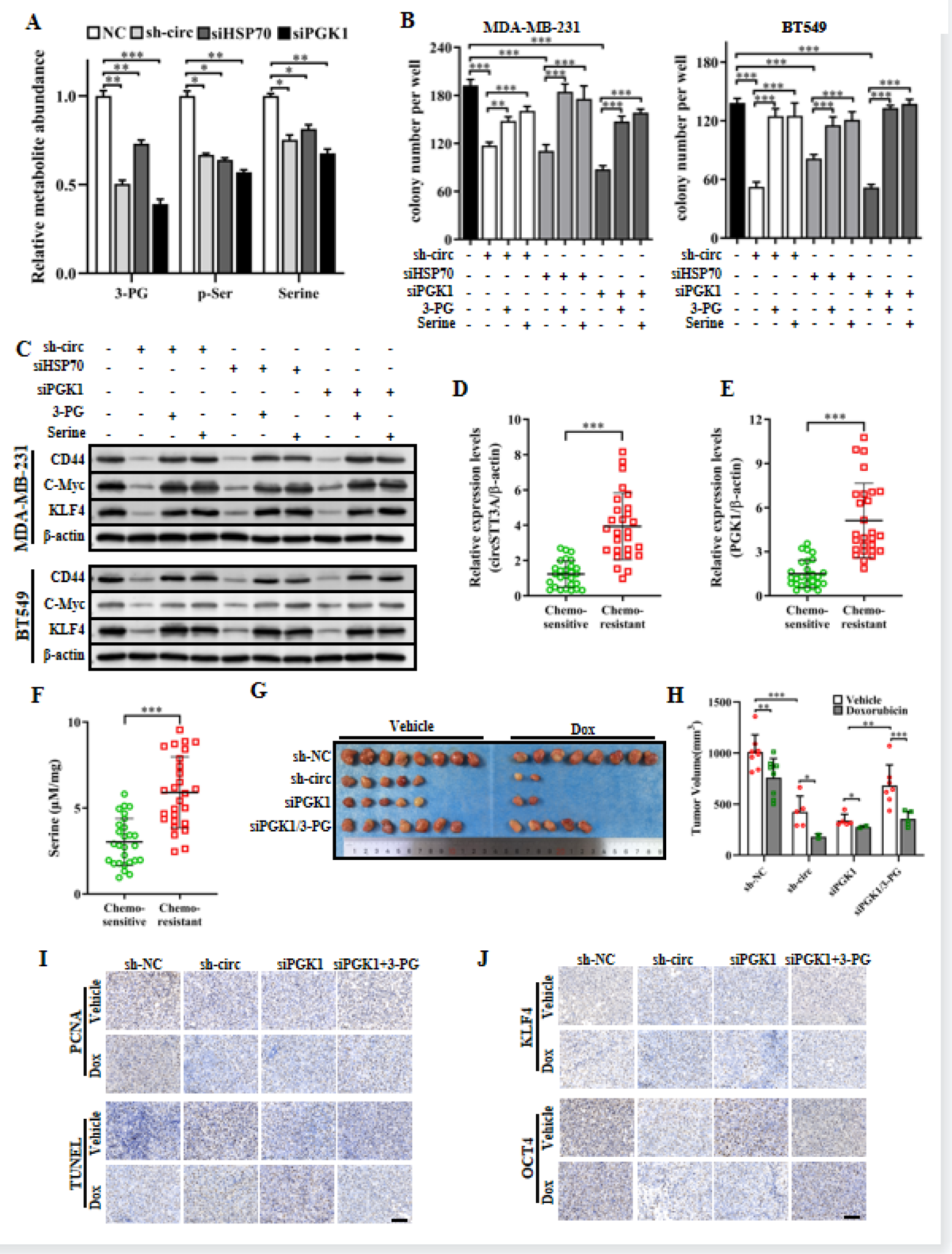
Enhanced metabolites related to the serine pathway endows BC with stemness maintenance and chemo-resistance in vivo and in vitro. **a.** Cellular 3-phosphoglycerate (3-PG), phosphoserine (pSer) and serine concentrations were measured by HPLC-MS in circSTT3A-knockdown, HSP70-knockdown and PGK1 knockdown MDA-MB-231 derived spheres and control spheres under hypoxia. **b-c**. Colony formation assays **(b)**, western blot analyses **(c)** of stemness-related proteins (CD44, c-Myc, and KLF4) in BC cells under indicated conditions (Scale bar, 200 µm). **d-f**. The expression of circSTT3A **(d)**, PGK1 **(e)** and serine **(f)** in the chemo-sensitive tissues (n=27) and chemo-resistant tissues (n=27) of BC patients. **g-h**. Representative pictures of xenograft tumors (n=8) **(g)** and xenograft tumor volume **(h). i-j**. Representative images showed PCNA IHC staining **(i)**, cell apoptotic status checked by TUNEL **(i)**, KLF4 and OCT4 levels **(j)** in each group of xenograft tumors (Scale bar, 100 µm). Data are presented as the mean±SD; *p<0.05, **p <0.01 and ***p < 0.001.

The consensus is that CSCs contribute to chemotherapy resistance in cancer progression, we further explored the related gene levels and serine concentration in BC patients with chemotherapy resistance. In BC tumor tissues, the levels of circSTT3A and PGK1 were significantly elevated in chemo-resistant tissues compared with chemo-sensitive tissues (Fig. 7d-7e). Meanwhile, as a metabolite for tumorigenesis promoter [22], the increased serine level was ascertained by serine detection kit in the chemo-resistant tissues of BC patients (Fig. 7f). To expand our findings, xenograft models of nude mice were employed. The mice injected with PGK1-silenced spheroid cells derived from hypoxic MDA-MB-231 attenuated tumor-initiating potentials and tumor growth, and exogenous addition of 3-PG could increase tumor initiating abilities and tumor growth in PGK1-functional loss tumors (Fig.7g-7h). In daily practice, Doxorubicin (Dox) was widely used as chemotherapeutic drug in BC patients. Thus, we tried to verify the effects of circSTT3A, PGK1 and 3-PG for BC tumor sensitivity to Dox (Fig.7g-7h). Loss of circSTT3A and PGK1 substantially suppressed tumor initiation and tumor growth, which dramatically increased tumor sensitivity to Doxorubicin treatment in mice; administrated tumor-burden mice injected PGK1-silenced spheroids with 3-PG could significantly reverse tumor initiation and tumor growth and increased tumor resistance to Doxorubicin. Correspondingly, the tumor growth and sensitivity to Doxorubicin in circSTT3A- and PGK1-deficent tumor were further assessed by PCNA and TUNEL staining (Fig. 7i) and stemness-related marker expressions of KLF4 and OCT4 by IHC staining (Fig. 7j). Taken together, these data denote that hypoxia-induced circSTT3A promotes 3-PG accumulation and serine synthesis via enhanced PGK1, which augments BCSCs stemness maintenance and chemotherapy resistance.

## Discussion

Hypoxia is one of the most important characteristics of solid tumors. A large number of experimental data in vitro and in vivo have shown that hypoxia coordinates the malignant phenotype of cancer cells by activating multiple oncogenic signaling pathways [23]. These pathways involved in metabolic reprogramming, angiogenesis, tissue remodeling, stemness, and immune regulation [24]. Evolving evidence supports that BC exists the plasticity between the breast cancer stem cells (BCSC) and non-BCSC compartments [25]. The formation and maintenance of BCSC are the main factors of recurrence and drug tolerance [26]. Studies have shown that under hypoxic conditions, the CSC formation efficiency will significantly increase, and this process is regulated by hypoxia-inducible factors [27]. However, the molecular mechanism of BCSC formation and stemness maintenance under hypoxic conditions is still unclear. We screened circSTT3A with abnormally elevated expression in BC cells under hypoxic conditions, and its transcription was regulated by HIF-1α, which could provide BC cells with greater flexibility to adapt to hypoxic environmental conditions. In addition, circSTT3A played an important role in relating the 3-PG accumulation and the serine synthesis, and drove the BCSC formation in hypoxia environment and enhances tumorigenesis in vivo. These findings suggested that hypoxia-induced circSTT3A was a key molecule in BCSCs plasticity, stemness and BC initiation.

CircRNAs, a circular RNA formed by back splicing, are considered a relatively conservative and stable RNA structure [28]. It is now appreciated that circRNAs could be used as biomarker and affect the phenotypic characteristics of cancer cells during tumor development [29–31]. A number of mechanisms are involved in the regulation of circRNAs expression, including hypoxic conditions. It has been reported that the expression of circWSB1 is regulated by HIF-1α under hypoxia, and could be used as a biomarker for BC [12]. In our study, clinical evidence indicated that HIF-1ɑ-induced circSTT3A was associated with the progression of BC patients. We demonstrated that circSTT3A had great diagnostic efficacy to sensitively distinguish BC. Moreover, the high expression of circSTT3A was an unfavorable factor with poor survival time in BC patients, and had a significant correlation with the pathological grade and T grade of BC. Compared to the primary BC cells with low level of circSTT3A, these cells with high expression of circSTT3A had higher sphere formation ability of BCSCs, more proportion in CD44^+^/CD24^−/low^ cell population, and stronger tumorigenesis. These results suggested that circSTT3A was involved in the stemness maintenance and tumorigenesis of BC, which may provide BC cells with more flexibility to adapt to hypoxia.

As a molecular chaperone, heat shock protein 70 (HSP70) plays a “seesaw” role in protein synthesis, folding, transportation, assembly, and degradation - balancing protein homeostasis from the perspective of post-translational processing [32, 33]. HSP70 is an accomplice in cancer progression and is potential as anti-cancer therapeutic targets. Despite tremendous efforts to explore the mechanism of HSP70, there remains necessity to understand the functional roles of HSP70 in cancer. HSP70 protein contains two important functional domains: N-terminal nucleotide-binding domain (NBD) and C-terminal substrate-binding domain (SBD) [34]. NBD can bind ATP and proteins to control SBD performing cellular functions of Hsp70 protein, thus some known Hsp70 inhibitors are designed to interfere with NBD [35]. Through mass spectrometry, RNA pull-down and RIP analyses, we provided the first evidence that circSTT3A directly interacts with HSP70 NBD and further stabilizes PGK1 protein via HSP70 SBD. Our immunopreciptation data showed circSTT3A directly binds with HSP70 NBD while artificial alteration of circSTT3A expression was unable to change the RNA transcription and protein level of HSP70 in BC cells. Meanwhile, high circSTT3A expression concentrated the interaction between HSP70 and PGK1, and further decreased the ubiquitination of PGK1. Briefly, circSTT3A as a non-code RNA, should modulate the interaction of HSP70 and PGK1, which provided a new mechanistic insight into the interconnected manner of HSP70 and downstream proteins in hypoxic BC cells.

Abnormally activated glycolysis is an emerging hallmark of many human cancers, and cancer cells acquire energy from glycolysis to promote growth, survival and proliferation [36]. Under hypoxic conditions, based on the intermediates from glycolysis, HIFs regulate transcription of enzymes to decrease the generation of acetyl CoA for entry into the tricarboxylic acid (TCA) cycle, and increase conversion of glucose to serine [37]. PGK1, the first ATP-generating enzyme in the glycolytic process, catalyzes 1,3-diphosphoglycerate (1,3-BPG) into 3-PG [38], which was found to be upregulated in BCSCs and some cancers of specific organs [39–41]. Blocking PGK1 led to decreased 3-PG concentration and inhibited tumor growth in pancreatic cancer [42]. It has been documented that 3-PG as the substrates is shunted to the de novo synthesis of serine, and then active serine synthesis signaling helps tumor stem cells adapt to hypoxia for cell survival [43]. Here, we unveiled that circSTT3A-HSP70 mediated ubiquitination changes of PGK1 acts an essential role for PGK1 stability, which contributes to accumulation of 3-PG and increase of serine synthesis for BC cell proliferation and BCSC formation under hypoxic microenvironment. Furthermore, the current work demonstrated that the enhanced circSTT3A, PGK1 and increased serine are positively correlated with BC patient chemo-resistance, suggesting that circSTT3A induced metabolism reprogramming is a crucial factor for tumor malignancy.

In summary, our works highlighted that HIF-1α-induced circSTT3A interacts with HS070 to increase PGK1 protein stability via modification of ubiquitination, which promotes accumulated 3-PG into serine synthesis to increase BCSC enrichment under hypoxic conditions. CircSTT3A is a potential biomarker in BC, and targeting circSTT3A and circSTT3A/HSP70/PGK1 combined with chemotherapy provides a new therapeutic strategy for breast cancer.

## Conclusions

In summary, we highlight HIF-1α-induced circSTT3A can interact with HS070 to increase PGK1 protein stability, which could regulate accumulation 3-PG into serine synthesis to increase BCSC enrichment under hypoxic conditions. These results demonstrate that circSTT3A is a potential biomarker in BC and circSTT3A/HSP70/PGK1 constitutes a positive feedback loop to BCSCs maintenance, tumor growth and drug resistance. Our works also suggest that circSTT3A is a potential biomarker in BC and provides a new insight into the regulation of BCSCs metabolic reprogramming by hypoxia and could clinical treatment strategy for BC.

### Abbreviations

BCSCs: breast cancer stem cells
BC: breast cancer
circRNA: circular RNA
HIF1α: hypoxia inducible factor 1 alpha
Hsp70: heat shock protein 70
PGK1: phosphoglycerate kinase 1
3-PG: 3-phosphoglycerate
1,3-BPG: 1,3-diphosphoglycerate
SSP: serine synthesis pathway
FISH: Fluorescence insitu hybridization
RT-PCR: Quantitative reverse transcription polymerase chain reaction
shRNA: Short hairpin RNA
IHC: Immunohistochemistry
ChIP: Chromatin immunoprecipitation
RIP: RNA immunoprecipitation
IF: Immunofuorescence
siRNA: Small interfering RNA
Co-IP: Co-immunoprecipitation

## Ethics approval and consent to participate

All human tumor tissue samples were collected in accordance with national and institutional ethical guidelines.

## Consent for publication

All authors approved the manuscript for submission and consented for publication.

## Availability of data and materials

The datasets supporting the conclusions of this article are included within the article and its supplementary files.

## Author contributions

Y.S. and M.L. designed the study. M.X., S.Y., and M.L wrote the manuscript. M.X. S.C., R.T., and X.Z. conducted the experiments. Y.Q., L.Y., Y.Q., and T.J. analyzed the data. S.W., and Y.G were responsible for clinical sample collection. S.Y., and M.L revised this manuscript.

## Acknowledgements

We thank the Department of Endocrine and Breast Surgery, The First Afliated Hospital of Chongqing Medical University for their helpful in collecting tumor samples and related anonymous clinical data.

